# Spatial landscape of pretransplant liver disease and post-transplant immune outcomes in children

**DOI:** 10.64898/2026.09.28.754819

**Authors:** Chris Kang, Mylarappa Ningappa, Brandon W Higgs, Catherine Gestrich, George Mazariegos, Chethan Ashokkumar, Lori Schmitt, Sarangarajan Ranganathan, Shankar Subramaniam, Rakesh Sindhi

## Abstract

Defining molecular patterns underlying pediatric end-stage liver disease and its treatment with liver transplantation requires rich datasets spanning a multitude of diagnoses and transplant outcomes. Here, we describe the spatial molecular landscape of approximately five million liver cell profiles from 98 liver tissue samples. The samples span donor liver, used here as a reference tissue, and end-stage diseases including biliary atresia, progressive familial intrahepatic cholestasis, hepatoblastoma, acute liver failure, and gestational alloimmune liver disease. Post-transplant liver allograft biopsies include those without and with T-cell-mediated and antibody-mediated rejection, mixed pathology, and outcome groups from immunosuppression withdrawal studies. Samples were profiled using the CosMx Human Universal Cell Characterization RNA Panel (1K) and the CosMx Human 6K Discovery Panel (6K). Preliminary exploratory analyses include cellular composition and gene abundance and variability across clinical cohorts. A twelve-gene signature comprising the genes with the highest variance implicates acute-phase response genes (e.g., *SAA1*) and profibrotic genes (e.g., *COL1A1*) as a shared link across these otherwise distinct conditions. This injury-repair-remodeling gene network warrants further investigation through higher-resolution, zone-specific spatial mapping to help clarify disease pathogenesis.

## Introduction

Pediatric end-stage liver disease has distinct genetic and developmental origins, and liver transplantation (LT), which is curative for these conditions, engages a maturing immune system against this background, creating a combination of factors unique to children. How these factors interact and their contribution to disease and outcomes after LT remain poorly defined. Detailed spatial mapping of this kind may help clarify disease pathogenesis, in the hope of enabling earlier, better-informed interventions that arrest progression to LT and improve post-transplant outcomes.

Previous single-cell liver atlases have characterized cell populations, age-related expression and fibrotic niches [1–4], while spatially resolved studies have mapped their organization within liver tissue [5, 6]. However, these studies have not combined single-cell spatial transcriptomics at scale with broad coverage of pediatric liver disease and transplantation. We have previously examined expression networks in pediatric transplant rejection [7], molecular pathways in biliary atresia (BA) [8, 9] and hepatoblastoma (HB) [10], and spatial expression patterns in rejection [11].

Here we extend our previous work using spatial molecular imaging [12] of nearly 5 million cells from 98 liver tissue samples spanning distinct biological processes. End-stage disease arising from structural defects that progress to cholestatic cirrhosis includes BA, the single largest indication for LT in children, and progressive familial intrahepatic cholestasis (PFIC) [9, 13, 14]. HB, an unresectable embryonal cancer, is the largest malignant indication for LT in children [15]. Acute liver failure (ALF) and gestational alloimmune liver disease (GALD) add a further immune-mediated dimension to pretransplant disease that may have parallels in the transplanted liver, for example, T-cell-mediated rejection (TCMR) and antibody-mediated rejection (ABMR), and tolerance [16, 17]. Ischemia-reperfusion injury, which begins with reperfusion of an otherwise normal, cold-preserved liver allograft, provides a bridge between the normal liver, post-transplant and pre-transplant pathology. This disease spectrum does not yet include infection or metabolic liver disease, but can serve as a reference to evaluate these and other conditions.

## Methods

Samples were profiled by CosMx spatial molecular imaging [12], using the 1K and 6K panels. We performed an early exploratory analysis to characterize expression patterns across the dataset, examining similarities among tissue samples and genes with relatively consistent or variable abundance within and between clinical cohorts. For the 6K panel, we annotated cells using InSituType [18] with reference profiles derived from cell clusters and marker-enriched populations. Annotations were assessed through internal marker and subject checks, available protein stains, and comparison with a separate marker-score method. Core assignment markers and additional checking markers are listed in Supplementary Table S1.

Cellular analyses included all 6K panel cells from 93 samples and 60 subjects across all available cohorts. Cell-type fractions and shares of each gene’s counts were calculated per sample within each acquisition, then averaged within subjects and across subjects in each cohort. Gene-count shares excluded subjects with no counts for that gene. Within-type abundance used log-transformed counts per million (CPM) normalized to the cell type’s total non-control counts. Marker abundance and detection fractions were summarized by medians within subjects, followed by cohort medians for abundance and means for detection.

To visualize similarities among cells, principal component analysis (PCA) and uniform manifold approximation and projection (UMAP) were applied separately to each RNA panel using 5,000 cells per acquisition (120,000 6K cells and 15,000 1K cells), retaining 2,000 of the 6K targets and all 998 available 1K targets from the respective panels. Genes were ranked by within-acquisition mean-bin-normalized dispersion. We compared log normalization with analytic Pearson residuals [19] using the same cells and genes. We inspected the 50-component scree curves and compared UMAPs using 20 and 40 components. The displayed UMAP uses Pearson residuals and 40 components. We also evaluated Harmony for acquisition adjustment but retained the unadjusted UMAP [20].

RNA transcript counts were summed per sample, normalized to CPM using non-control targets, and transformed as log_2_ (CPM +1). Within each cohort, abundance was summarized by taking the median across each subject’s samples, then across subjects, giving subjects equal weight [21]. Exploratory minimal analysis thresholds of three subjects and two acquisitions per cohort were chosen to limit dependence on one or two subjects or a single acquisition, as each cohort contributed equally to the gene ranking. 73 6K samples from 45 subjects across ten cohorts met these requirements for gene selection. Final transcript targets were ranked by variance of these ten cohort medians. Our threshold selection was validated by assessing sensitivity to cohort composition by repeating the ranking across all 18 cohorts and then omitting each cohort in turn.

To identify genes with consistent expression across cohorts, we computed log_2_ ratios of target counts to the sample’s 95th-percentile negative-probe count and summarized these within subjects and cohorts. Genes required positive ratios in at least 80% of subjects in every cohort, a cohort-median abundance range of at most one log-transformed CPM unit, and a within-cohort subject interquartile range of at most one unit. Passing genes were ranked by their maximum within-cohort interquartile range.

## Results

### Samples and clinical cohorts

The study cohort includes 93 samples from 60 subjects across 24 acquisitions and 18 clinical categories profiled using the 6K panel, with five additional samples profiled with the 1K panel for a total of nearly 5 million cells (Table 1). One subject contributed samples to both panels. Figure 1 summarizes the cohort coverage and spatial distribution of annotated cell types in selected biopsy samples.

**Table 1:** RNA measurement coverage.

|  | 6K panel | 1K panel | Combined |
| --- | --- | --- | --- |
| Studies | 7 | 2 | 9 |
| Samples | 93 | 5 | 98 |
| Subjects | 60 | 3 | 62 |
| Cell profiles | 4,552,119 | 402,648 | 4,954,767 |

**Figure 1:**
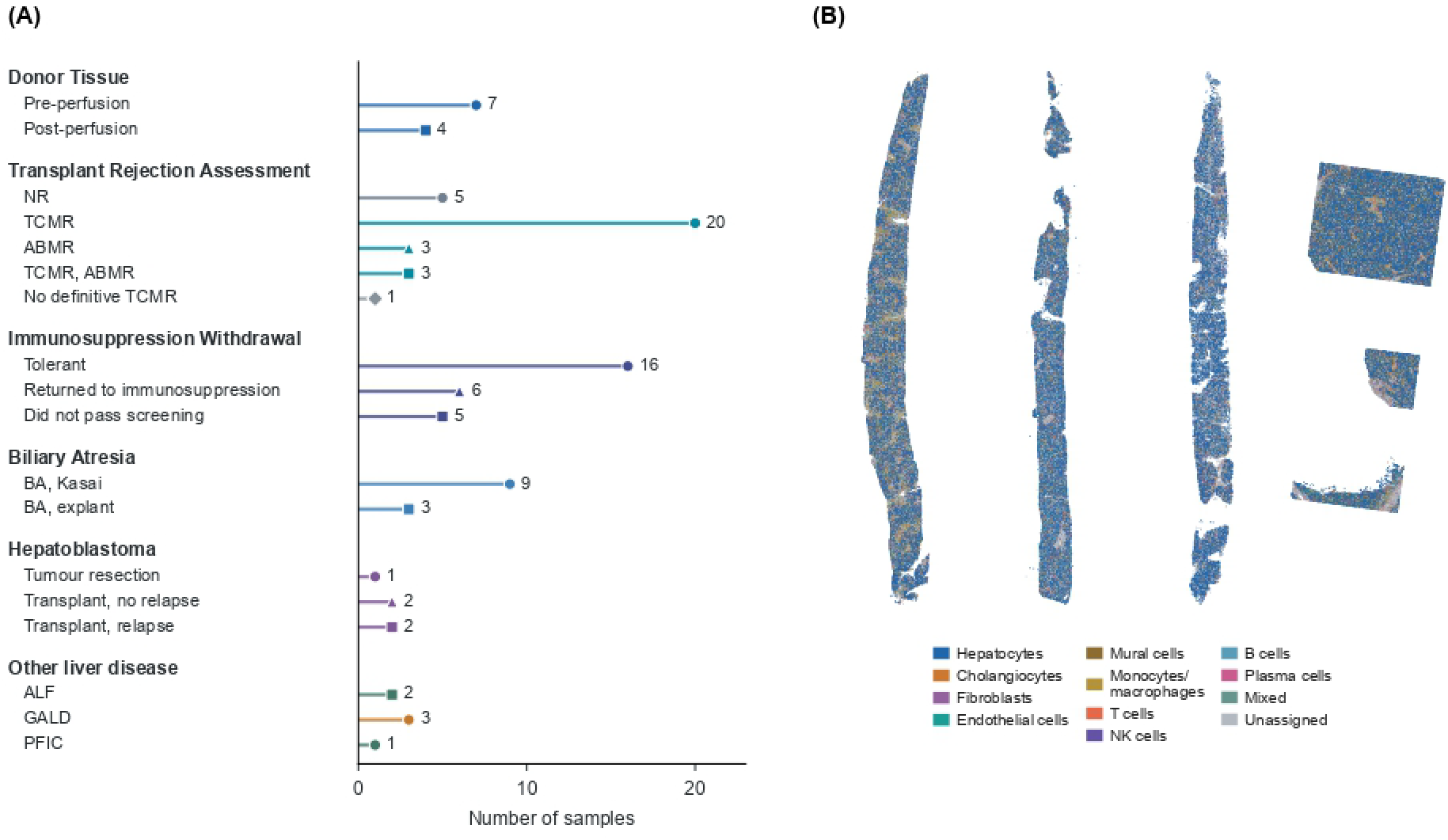
Cohort coverage and spatial sample maps. **(A)** Sample counts across 18 clinical categories, comprising 93 samples profiled using the CosMx Human 6K Discovery Panel. **(B)** Representative tissue samples from TCMR, immunosuppression withdrawal (tolerant), biliary atresia, and hepatoblastoma cohorts, from left to right. Each point represents a cell, colored by its annotated cell type.

The 18 cohorts are ordered in the various figures to reflect baseline (donor liver before and after perfusion), allograft liver with immunological outcomes, and native liver disease and its progressive states where possible. Hepatocyte proportions are relatively stable between paired donor liver samples before and after reperfusion. The next eight cohorts represent liver allograft biopsies with immunological findings: non-rejector (NR) biopsies despite elevated liver function tests, TCMR alone, ABMR, TCMR with ABMR [22], and three cohorts from our center’s participation in the multi-center Immunosuppression Withdrawal for Stable Pediatric Liver Transplant Recipients (iWITH) study [23]. In this study, biopsies were obtained at screening and end of study and in between when immunosuppression (IS) withdrawal was complete. The cohorts include those from subjects in whom IS withdrawal was successful over consecutive biopsies (tolerant), those representing failure of IS withdrawal with three biopsies from each of two subjects, and biopsies indicating histological inflammation at onset representing screen failure. These are followed by a range of native liver diseases: BA at the time of Kasai portoenterostomy; BA explanted at LT [13], where increasing fibrosis reflects cirrhotic explanted tissue; HB resected successfully, and HB that spread locally to replace hepatocytes, became unresectable, and required LT [15], after which one subgroup relapsed and the other did not; ALF, showing marked hepatocyte loss [16]; GALD, in which maternal antibodies deplete hepatocytes and cause liver failure requiring LT shortly after birth [17]; and PFIC explanted at LT [14]. Relative cell proportions in the smallest cohorts, such as the single sample with no definitive TCMR and the three with mixed TCMR/ABMR features, can be distorted because of sampling error.

### Data quality

Eleven samples in two acquisitions had elevated negative-probe measurements. However, elevated negative-probe measurements were not used as an exclusion criterion, and these samples were retained in all cellular analyses. For cells profiled with the 6K panel, the absolute Spearman correlation between the first principal component (PC1) and detected-gene count decreased from 0.85 with log normalization to 0.10 with Pearson residuals, and we used Pearson residuals for all displayed UMAPs. Acquisition adjustment (Harmony) increased mixing between acquisitions but also reduced agreement within clinical cohorts and marker groups, so we retained the unadjusted embedding as primary and Harmony as a sensitivity analysis. UMAP neighborhoods varied with the number of input components and were not interpreted as stable cell states. Acquisition and tissue context were confounded in this cohort, so the cell-type composition and gene-attribution results below rely on per-cell marker-based classification rather than on UMAP cluster boundaries.

### Cell composition

We first examined cellular composition and then asked which genes vary most across clinical cohorts and which show relatively consistent expression.

The 6K collection included hepatocytes, cholangiocytes, fibroblasts and endothelial cells, together with mural cells, monocytes/macrophages, T cells, natural killer (NK) cells, B cells and plasma cells (Figure 2). Hepatocytes had the largest subject-balanced cell fraction among identified types in 15 of 18 cohorts (Figure 2B). Mixed and unassigned cells accounted for 34.5% of cells from the 93 6K samples and remained in the quantitative summaries. Finer immune states were not resolved.

**Figure 2:**
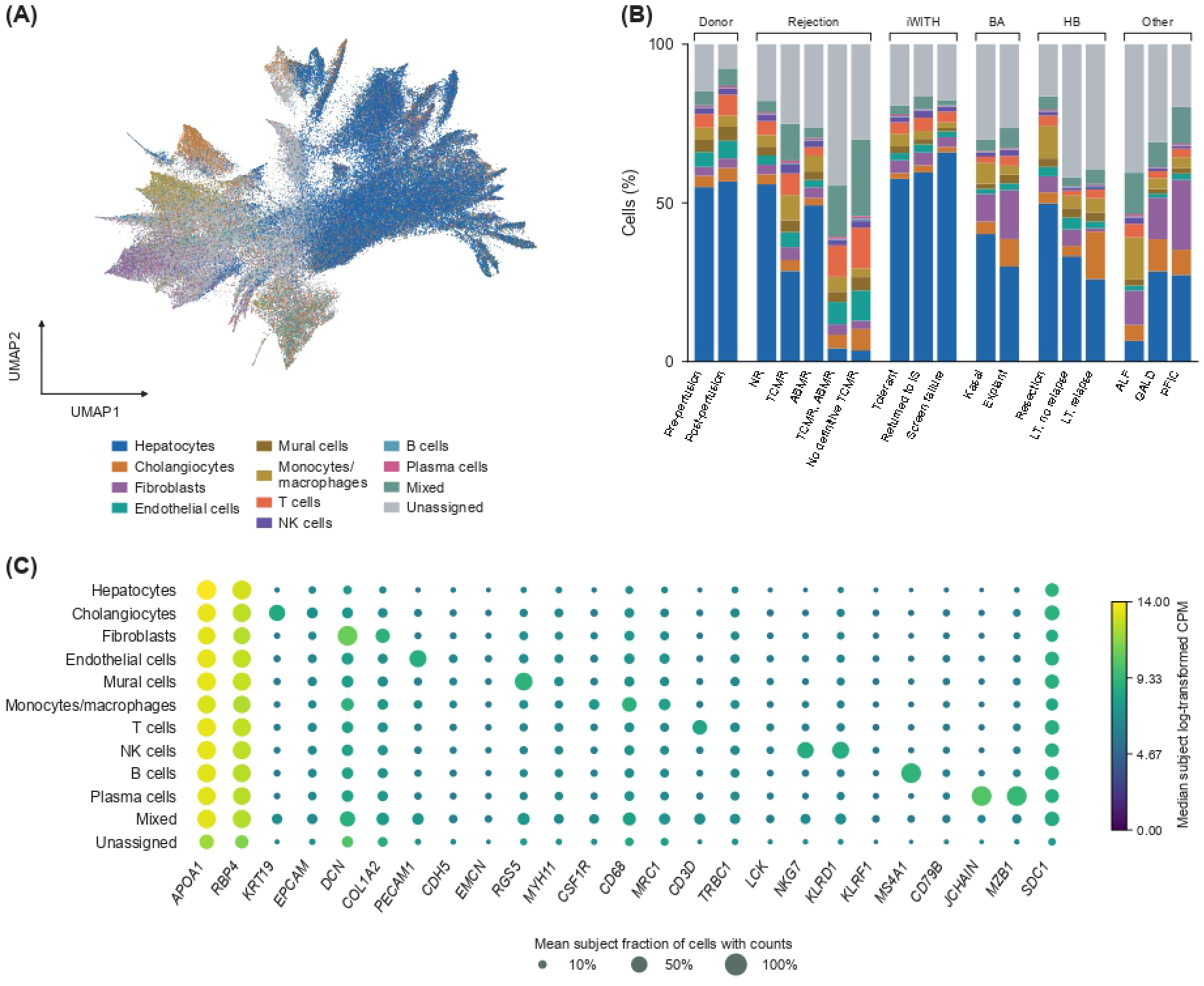
Cell types and transcript expression. **(A)** The 6K UMAP constructed from 2,000 selected genes, colored by InSituType cell types. The map contains 120,000 acquisition-balanced cells. Individual cell type UMAPs are shown in Supplementary Figure S1. **(B)** Subject-balanced cell-type proportions across 18 clinical cohorts, using 4,552,119 cells from 93 samples and 60 subjects. **(C)** Expression of 25 liver and immune markers across these groups. Dot area represents the mean subject-level fraction of cells with counts for the marker.

As described in the previous section, samples of native liver disease that progressed to LT, such as BA and PFIC explants, demonstrated increased proportions of fibroblasts consistent with cirrhosis. Conditions with acute loss of liver function, such as ALF and GALD, showed marked loss of hepatocyte fractions, as did locally invasive and unresectable HB compared with localized and resectable HB. Among the eight additional biopsy cohorts representing allograft liver with various immunological outcomes, the single sample with no definitive TCMR and the three with mixed TCMR/ABMR features showed markedly low hepatocyte fractions, approaching those seen in ALF. Because rejection is rarely associated with loss of synthetic function as occurs in ALF and GALD, this finding more likely reflects sampling and sample-processing variability than true hepatocyte loss.

### Transcript variation across cohorts

Initial gene ranking was restricted to ten cohorts with at least three subjects and two acquisitions each to limit sampling variability under equal cohort weighting. Across these ten cohorts, *SAA1* and *COL1A1* showed the greatest variance in cohort median abundance (Figure 3A). Rankings were then recomputed across all 18 cohorts to assess sensitivity to cohort composition. Eight of the twelve leading targets remained among the top twelve, with *SAA1* ranked first and *COL1A1* eighth. Omitting individual cohorts from this expanded analysis gave ranks of 1–2 and 4–16, respectively, indicating sensitivity to cohort composition.

**Figure 3:**
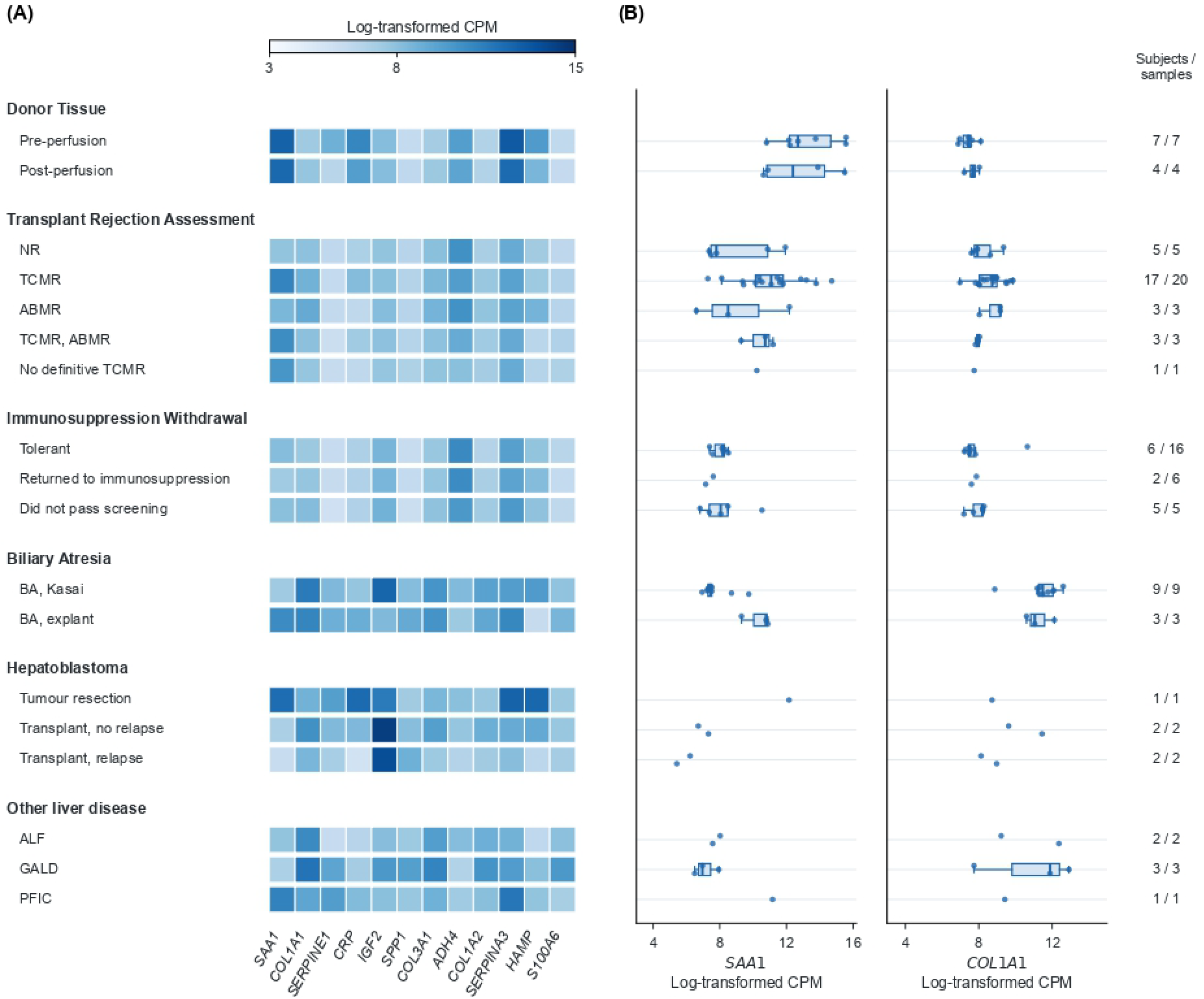
Gene abundance and variation across cohorts. **(A)** Expression across all 18 clinical cohorts for twelve targets selected by the greatest variance in subject-balanced medians across the ten cohorts meeting selection threshold criteria. **(B)** Subject-level abundance of *SAA1* and *COL1A1*, showing the two highest-variance targets in the ten-cohort selection.

We therefore retained *SAA1*, the highest-ranked acute-phase gene, and *COL1A1*, a prototypical collagen gene, as representative targets for examining acute-phase and fibrotic responses. Donor tissue before perfusion had higher median *SAA1* and lower median *COL1A1* abundance than GALD (Figure 3B). Among low-variation targets across the ten cohorts used for gene selection (73 samples; 45 subjects), *RAD23B* had a cohort median range of 0.39 log-transformed CPM units and a maximum within-cohort interquartile range of 0.28. Figure 3 shows the 12 selected genes across all 18 cohorts. Gene selection and the low-variation summary use the ten-cohort comparison.

To relate cohort-level variation to cellular composition, we examined each cell type’s share of gene counts across the 6K samples (Figure 4). Hepatocytes contributed the largest share of *SAA1* counts in 15 of 18 cohorts, or 14 without the core-marker requirement. Hepatocytes also contributed the largest share of *HP* counts in most cohorts.

**Figure 4:**
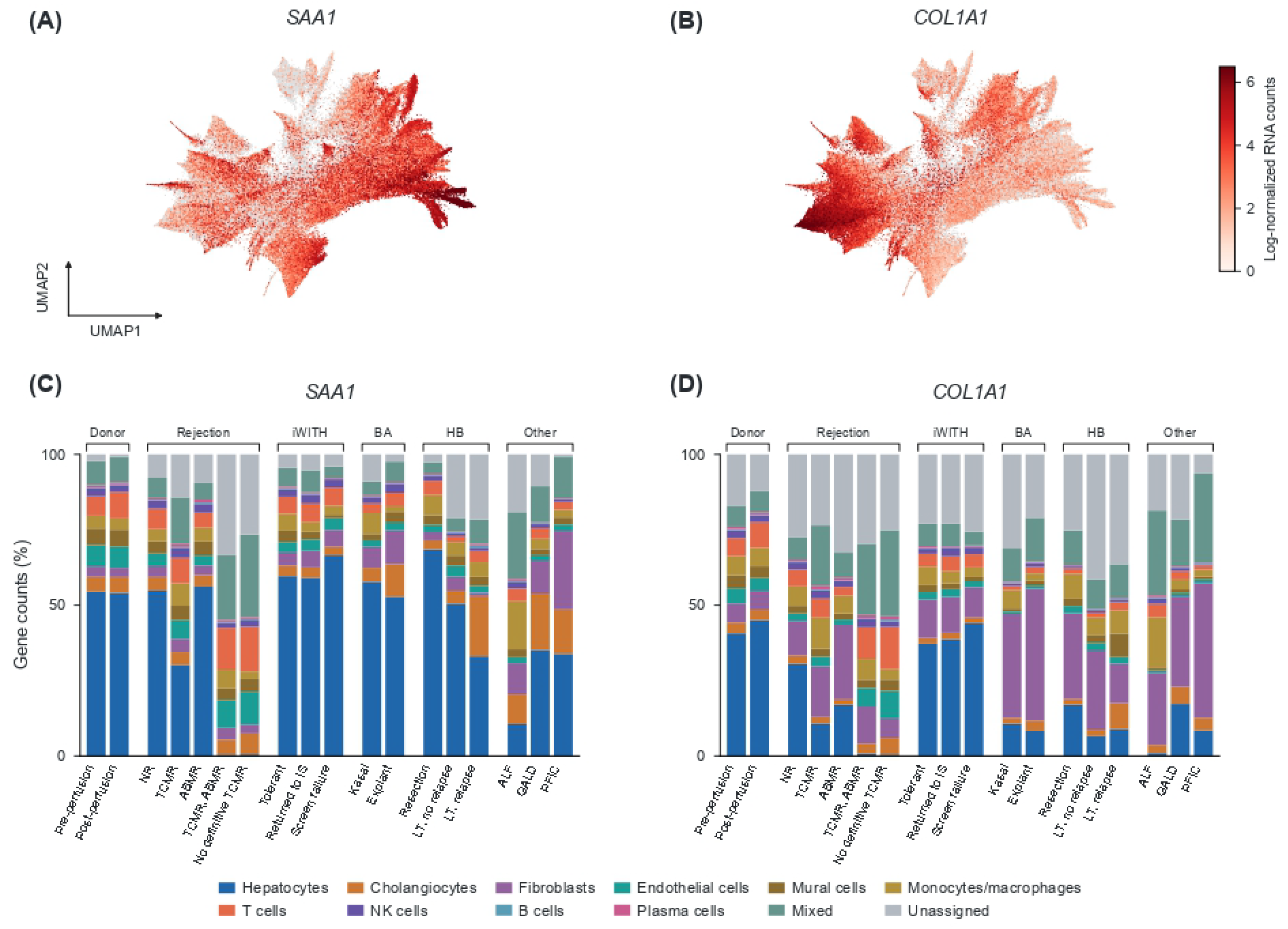
Cellular gene expression and transcript contributions. **(A,B)** *SAA1* and *COL1A1* abundance over the same 6K Pearson-residual UMAP of 120,000 cells. Color shows natural-log-transformed counts per 10,000 RNA counts, on a shared scale from 0 to 6.5; higher values saturate. Gray cells have zero counts for the displayed gene. **(C,D)** Subject-balanced fractions of each gene’s observed counts assigned to the cell types in Figure 2, across all 18 clinical cohorts.

Among the named cell types, fibroblasts had the highest within-group normalized *COL1A1* abundance in all 18 cohorts, including the assignment sensitivity. Fibroblasts contributed the largest share of total *COL1A1* counts in five cohorts, while hepatocytes contributed the largest share in six; mixed or unassigned cells led in the remaining seven (Figure 4D).

### Cellular and transcriptional changes between paired outcomes

To examine how cell-type distribution and gene expression changes demonstrate differences between related clinical outcomes, we analyzed two illustrative pairs: NR versus TCMR, and ALF versus GALD (Figure 5). Compared with biopsies without rejection (NR), TCMR biopsies showed a broader spatial distribution of non-hepatocyte cell types, with a corresponding drop in subject-balanced hepatocyte fraction (from approximately 55% to 30%) and an expansion of immune and mixed populations (Figure 5C). In the ALF/GALD pair, both conditions showed substantially lower hepatocyte fractions than either rejection group, consistent with acute loss of hepatocyte mass in both. Additionally, GALD showed larger fibroblast and cholangiocyte fractions than ALF.

**Figure 5:**
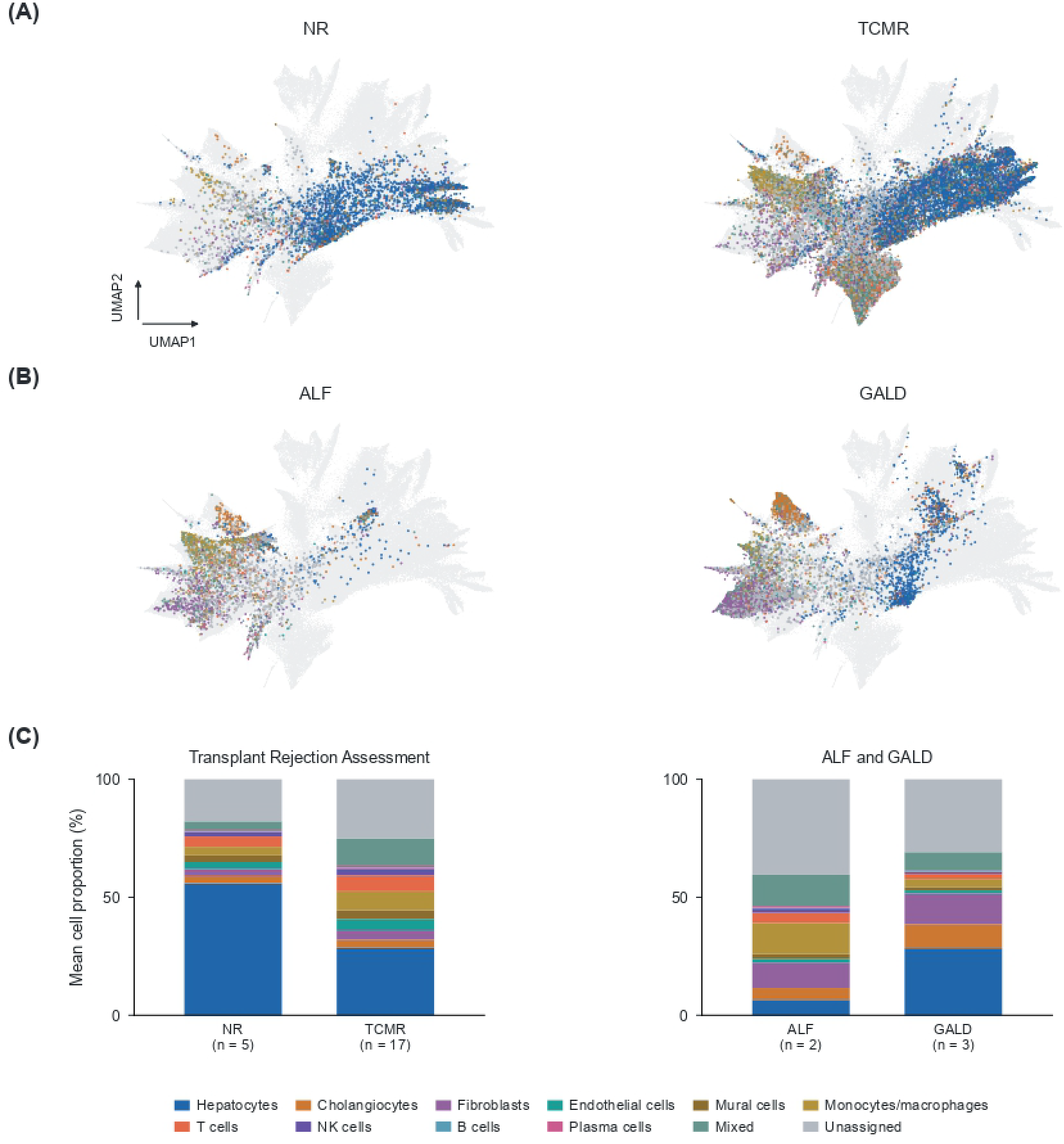
Cohort-specific cell distributions in UMAP space of allograft and native liver disease states. Cells from paired clinical subcategories are overlaid on the full 6K UMAP (gray, as in Figure 2A) and colored by annotated cell type. **(A)** NR (left, *n* = 5) and TCMR (right, *n* = 17). **(B)** ALF (left, *n* = 2) and GALD (right, *n* = 3). **(C)** Subject-balanced cell-type proportions for each pair shown in panels A and B.

These compositional differences were accompanied by distinct combinations of six of the 12 genes with the greatest variance and their cellular origins in each pair (Figure 6). Within the rejection pair, *SAA1* was the dominant discriminating signal, and was elevated in TCMR relative to no rejection across nearly every annotated cell type, with a smaller parallel increase in *CRP*. Collagen transcripts (*COL1A1, COL1A2, COL3A1*) were only modestly higher in TCMR. Within the ALF/GALD pair, the pattern was reversed: collagen transcripts were consistently and substantially higher across nearly all cell types in GALD relative to ALF, while *SAA1* and *HP* were lower in GALD than ALF.

**Figure 6:**
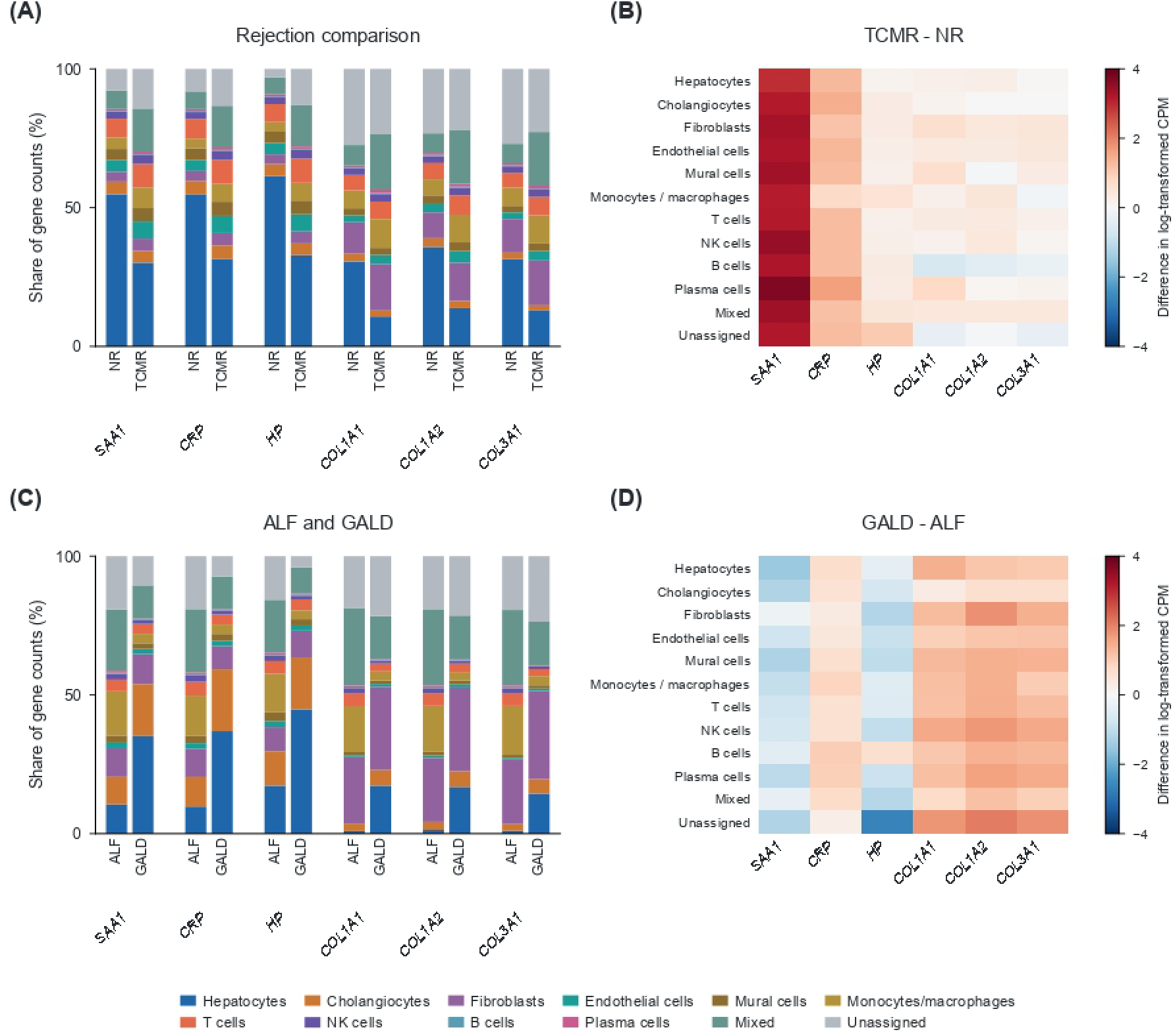
Cell-type contributions to acute-phase and fibrotic gene expression across paired clinical subcategories. Six targets (*SAA1, CRP, HP, COL1A1, COL1A2, COL3A1*) are shown for the same two cohort pairs as Figure 5. **(A,C)** Subject-balanced share of each gene’s counts (%) contributed by each annotated cell type, for **(A)** NR versus TCMR, and **(C)** ALF versus GALD. **(B,D)** Difference in cell-type-specific, log_2_-transformed CPM abundance for the same six targets, comparing(**B**)TCMR minus NR, and **(D)** GALD minus ALF. Red indicates higher abundance in the first-named group of each comparison; blue indicates lower abundance.

Together, these two paired comparisons suggest that the acute-phase and fibrotic signatures identified across the full cohort (Figure 3) are not uniformly co-elevated in every disease state, but instead segregate. Subcategories within the rejection pair are distinguished primarily by acute-phase/inflammatory genes, while the ALF/GALD pair is distinguished primarily by the accompanying fibrosis in GALD. This raises the possibility that the same shared molecular pathways consisting of acute-phase reactants and fibrotic genes is deployed differently, for example via different cell types, depending on the specific pre- or post-transplant process, rather than reflecting a single common injury pathway that behaves identically across all disease cohorts.

## Discussion

At its current stage, our whole-biopsy-based description of cellular composition and gene expression across a set of illustrative clinical cohorts that present most commonly for LT in childhood is descriptive, rather than a mechanistic or diagnostic study. With that caveat, three observations from this initial analysis are worth highlighting.

First, cell-type composition varied systematically with clinical cohort in ways that reflect disease progression toward LT. Using donor liver obtained before or shortly after cold preservation as a relatively normal reference point, we observed a marked loss of hepatocytes in conditions with acute loss of synthetic function, such as ALF and GALD, and an increase in fibroblasts in cirrhotic liver diseases such as BA and PFIC that progress to LT. At this stage of data accrual these cohort-specific findings are illustrative. Donor tissue, while the closest available reference to a normal liver in this cohort is itself procured under conditions of brain death or circulatory arrest and subjected to cold preservation and a perfectly healthy baseline may be challenging.

Second, the twelve genes with the greatest variance in abundance across biopsies and cohorts largely organize around two related categories: acute-phase reactants, including *SAA1, CRP, HP, SERPINA3, HAMP* and *S100A6*, and matrix and growth-associated genes linked to fibrosis, including *COL1A1, COL1A2, COL3A1, SERPINE1* and *IGF2*. A STRING protein-protein interaction network restricted to these twelve genes placed *IGF2* with the fibrotic set, connecting directly to *SERPINE1* and *COL1A1* but to none of the acute-phase genes, consistent with its reported role in hepatic stellate cell activation and fibrogenic signaling. *ADH4*, a hepatocyte metabolic enzyme, showed no direct network connection to either group and was not assigned to either category. In the liver, acute-phase and fibrotic gene programs mark injury, repair and remodeling, which are related processes that co-occur in response to insults of many kinds, whether immunological, ischemic, or obstructive [24]. Their recurrence as the dominant sources of variation across this diverse set of cohorts suggests shared molecular injury pathways underlying otherwise distinct pediatric liver diseases and post-transplant outcomes. This inference is consistent with the hepatic acute-phase role of serum amyloid A (SAA) and its predominant hepatocyte production [25], and with the role of collagen-producing mesenchymal cells in the liver fibrosis niche [4].

Third, resolving the relative contribution of cell types to these variable genes has the potential to distinguish closely related outcomes that are difficult to separate by cell composition or gene expression alone, or by histological criteria. This potential is illustrated in part by rejection versus no rejection and by ALF versus GALD (Figures 5 and 6). These two pairs of related outcomes separated along different injury-repair-remodeling axes within the shared signature: the rejection pair separated primarily along the acute-phase reactant gene axis, while the ALF/GALD pair separates primarily along the fibrotic gene axis, suggesting that the same underlying injury pathways are deployed differently and via different cell types, depending on the specific disease process. This discriminative power is likely to sharpen further once these signals are resolved by lobular zone, since many of these diagnoses carry well-characterized, zone-specific histological criteria. TCMR is a clear example. The Banff classification defines the diagnostic injury pattern of TCMR as T-cell-rich lymphocytic infiltration concentrated around cholangiocytes and endothelium in the portal and pericentral regions [22]. Thus, cell-type-specific expression of genes with high variance, further resolved by zonal distribution and anchored to the histological ground truth, is a natural next step for our future efforts.

Our current work has additional obvious limitations. Molecular image acquisition occurred over 24 separate runs, limiting our ability to fully distinguish technical batch effects from biological variation. However, based on cell type distribution, the data appear to be well balanced, and analyses are ongoing to address this limitation. Several of the illustrative cohorts, notably PFIC with one sample, indeterminate for TCMR with one sample, and the mixed TCMR, ABMR cohort with three samples, are small and could be confounded by sampling error whereby lobular architecture may not be represented accurately in these biopsies. Here, our findings should be considered exploratory. This cohort does not yet include infection or metabolic liver disease. Finally, this analysis does not yet incorporate lobular zonation, cell-cell interaction modeling, unsupervised clustering approaches, or further resolution of unassigned cell types, or unsupervised clustering approaches, each of which could further refine mechanisms underlying each disease state described here, in our ongoing follow-up analyses.

## Supplementary Information

## Abbreviations

ABMR: antibody-mediated rejection
ALF: acute liver failure
BA: biliary atresia
CPM: counts per million
GALD: gestational alloimmune liver disease
HB: hepatoblastoma
IS: immunosuppression
iWITH: Immunosuppression Withdrawal for Stable Pediatric Liver Transplant Recipients
LT: liver transplantation
NK: natural killer
NR: non-rejector
PCA: principal component analysis
PFIC: progressive familial intrahepatic cholestasis
RNA: ribonucleic acid
SAA: serum amyloid A
TCMR: T-cell-mediated rejection
UMAP: uniform manifold approximation and projection.

---

**Table S1:**
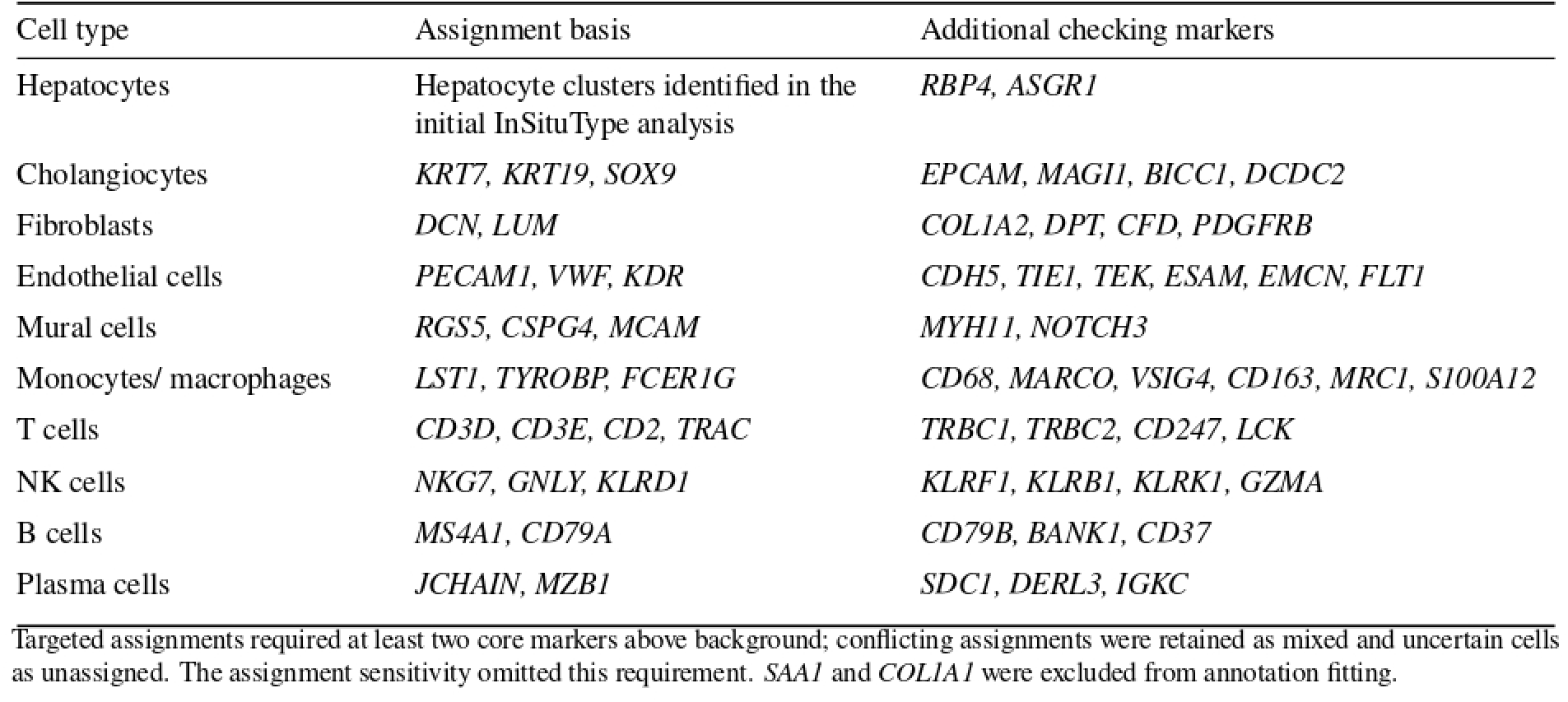
Cell-type assignment and checking markers.

**Figure S1:**
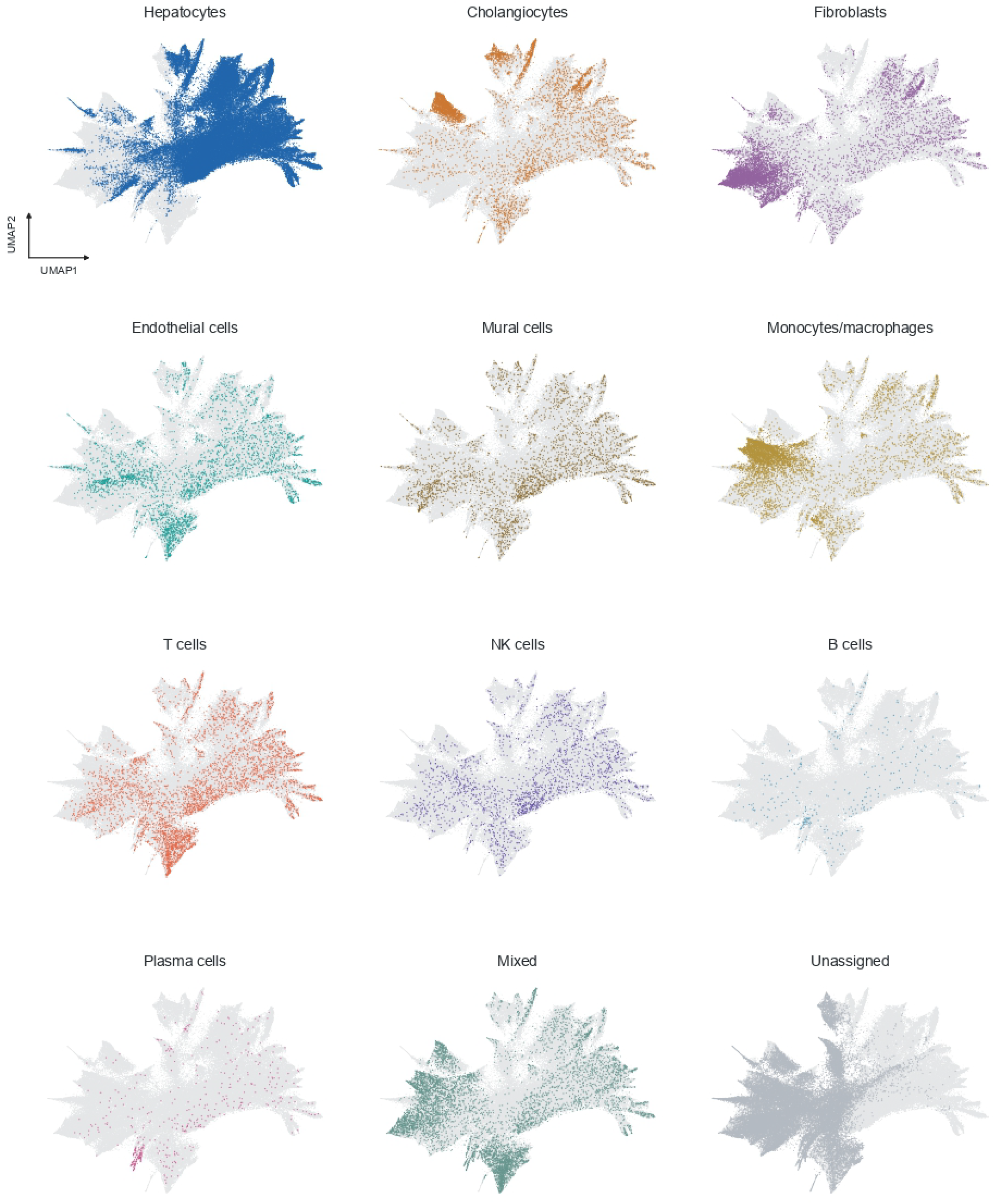
Cell-type distributions in UMAP space. Each panel highlights one cell type annotation category on the same UMAP shown in Figure 2A. All other cell types remain gray.

## References

[1] Nadim Aizarani, Antonio Saviano Sagar, et al. A human liver cell atlas reveals heterogeneity and epithelial progenitors. Nature, 572:199–204, 2019. doi: 10.1038/s41586-019-1373-2.

[2] Sonya A. MacParland, Jeff C. Liu, Xue-Zhong Ma, et al. Single cell RNA sequencing of human liver reveals distinct intrahepatic macrophage populations. Nature Communications, 9:4383, 2018. doi: 10.1038/s41467-018-06318-7.

[3] Rachel D. Edgar, Diana Nakib, Damra Camat, et al. Single-cell atlas of human pediatric liver reveals age-related hepatic gene signatures. Hepatology Communications, 9(11):e0813, 2025. doi: 10.1097/HC9.0000000000000813.

[4] Prakash Ramachandran, Ross Dobie, John R. Wilson-Kanamori, et al. Resolving the fibrotic niche of human liver cirrhosis at single-cell level. Nature, 575:512–518, 2019. doi: 10.1038/s41586-019-1631-3.

[5] Brianna R. Watson, Biplab Paul, Raza Ur Rahman, et al. Spatial transcriptomics of healthy and fibrotic human liver at single-cell resolution. Nature Communications, 16:319, 2025. doi: 10.1038/s41467-024-55325-4.

[6] Faizan Hasan, Rachel D. Edgar, Jawairia Atif, et al. High-resolution spatial transcriptomics of adult and pediatric human liver with Visium HD. Scientific Data, 2026. doi: 10.1038/s41597-026-07597-2. Published online 24 June 2026.

[7] Mylarappa Ningappa, Syed A. Rahman, Brandon W. Higgs, et al. A network-based approach to identify expression modules underlying rejection in pediatric liver transplantation. Cell Reports Medicine, 3(4):100605, 2022. doi: 10.1016/j.xcrm.2022.100605.

[8] Jun Min, Mylarappa Ningappa, Juhoon So, et al. Systems analysis of biliary atresia through integration of high-throughput biological data. Frontiers in Physiology, 11:966, 2020. doi: 10.3389/fphys.2020.00966.

[9] Joseph T. Glessner, Mylarappa B. Ningappa, Kim A. Ngo, et al. Biliary atresia is associated with polygenic susceptibility in ciliogenesis and planar polarity effector genes. Journal of Hepatology, 79(6):1385–1395, 2023. doi: 10.1016/j.jhep.2023.07.039.

[10] Sarangarajan Ranganathan, Mylarappa Ningappa, Chethan Ashokkumar, et al. Loss of EGFR–ASAP1 signaling in metastatic and unresectable hepatoblastoma. Scientific Reports, 6:38347, 2016. doi: 10.1038/srep38347.

[11] Mylarappa Ningappa, Chethan Ashokkumar, Divya Murali, et al. Liver transplant rejection features memory cell priming, spatially differentiated networks and novel drug targets. bioRxiv, 2025. doi: 10.1101/2025.02.04.635308. Preprint, posted 7 February 2025.

[12] Shanshan He, Ruchir Bhatt, Carl Brown, et al. High-plex imaging of RNA and proteins at subcellular resolution in fixed tissue by spatial molecular imaging. Nature Biotechnology, 40(12):1794–1806, 2022. doi: 10.1038/s41587-022-01483-z.

[13] J. B. Otte, J. de Ville de Goyet, R. Reding, et al. Sequential treatment of biliary atresia with Kasai portoenterostomy and liver transplantation: a review. Hepatology, 20(1 Pt 2):41S–48S, 1994. doi: 10.1016/0270-9139(94)90272-0.

[14] Mithat Gunaydin and Asudan Tugce Bozkurter Cil. Progressive familial intrahepatic cholestasis: diagnosis, management, and treatment. Hepatic Medicine: Evidence and Research, 10:95–104, 2018. doi: 10.2147/HMER.S137209.

[15] Ruy J. Cruz, Jr., Sarangarajan Ranganathan, George Mazariegos, et al. Analysis of national and singlecenter incidence and survival after liver transplantation for hepatoblastoma: new trends and future opportunities. Surgery, 153(2):150–159, 2013. doi: 10.1016/j.surg.2012.11.006.

[16] Catherine A. Chapin, T. Burn, T. Meijome, et al. Indeterminate pediatric acute liver failure is uniquely characterized by a CD103+ CD8+ T-cell infiltrate. Hepatology, 68(3):1087–1100, 2018. doi: 10.1002/hep.29901.

[17] Katie R. Conover, Samantha Saul, Catherine A. Chapin, et al. Defining a histologic scoring system for gestational alloimmune liver disease. The American Journal of Surgical Pathology, 49(12):1326–1333, 2025. doi: 10.1097/PAS.0000000000002454.

[18] P. Danaher, E. Zhao, Z. Yang, et al. InSituType: likelihood-based cell typing for single cell spatial transcriptomics. bioRxiv, 2022. doi: 10.1101/2022.10.19.512902. Preprint.

[19] Jan Lause, Philipp Berens, and Dmitry Kobak. Analytic Pearson residuals for normalization of single-cell RNA-seq UMI data. Genome Biology, 22:258, 2021. doi: 10.1186/s13059-021-02451-7.

[20] Ilya Korsunsky, Nghia Millard, Jean Fan, et al. Fast, sensitive and accurate integration of single-cell data with Harmony. Nature Methods, 16:1289–1296, 2019. doi: 10.1038/s41592-019-0619-0.

[21] Jordan W. Squair, Matthieu Gautier, Claudia Kathe, et al. Confronting false discoveries in single-cell differential expression. Nature Communications, 12:5692, 2021. doi: 10.1038/s41467-021-25960-2.

[22] A. J. Demetris, C. Bellamy, S. G. Hübscher, et al. 2016 comprehensive update of the Banff working group on liver allograft pathology: Introduction of antibody-mediated rejection. American Journal of Transplantation, 16(10):2816–2835, 2016. doi: 10.1111/ajt.13909.

[23] Sandy Feng, John C. Bucuvalas, George V. Mazariegos, et al. Efficacy and safety of immunosuppression withdrawal in pediatric liver transplant recipients: Moving toward personalized management. Hepatology, 73(5):1985–2004, 2021. doi: 10.1002/hep.31520.

[24] Antonella Pellicoro, Prakash Ramachandran, John P Iredale, and Jonathan A Fallowfield. Liver fibrosis and repair: immune regulation of wound healing in a solid organ. Nature Reviews Immunology, 14(3):181–194, 2014.

[25] S. V. Siegmund, M. Schlosser, F. A. Schildberg, et al. Serum amyloid a induces inflammation, proliferation and cell death in activated hepatic stellate cells. PLoS ONE, 11(3):e0150893, 2016. doi: 10.1371/journal.pone.0150893.

